# Phyllosphere bacterial communities in milkweeds: composition diverge as the season progresses, and links to cardenolides and arthropods depend on host identity

**DOI:** 10.64898/2026.09.21.753138

**Authors:** Danilo Ferreira Borges dos Santos, Victoria M. Pocius, Luana Bresciani, C. Guilherme Becker, Jared G. Ali, Francisco Dini-Andreote, Mônica F. Kersch-Becker

## Abstract

- Leaf bacterial communities shape plant defense and interactions with herbivores, yet how host filtering and stochastic processes assemble them remains unclear. We predicted that host identity, chemistry, and arthropods structure them, and that stochastic and deterministic contributions shift seasonally.
- Across one growing season we sampled leaf bacteria monthly and arthropods weekly on four Asclepias species (Apocynaceae: *A. curassavica*, *A. incarnata*, *A. syriaca*, *A. tuberosa*) spanning a cardenolide gradient. We sequenced 16S rRNA genes and quantified selection and dispersal contributions to turnover using null models.
- Host identity shaped bacterial richness and composition, with A. tuberosa hosting the richest communities, and both richness and phylogenetic diversity rose through the season. Early on, homogenizing dispersal made communities more similar among plants; by mid-season no single process dominated turnover, and by late season dispersal limitation prevailed. Homogeneous selection was episodic, not sustained, and arthropod associations were host-specific, negatively so on A. syriaca.
- Phyllosphere assembly thus shifts from convergence to divergence in one season: young leaves recruit from a shared pool delivered by wind, rain, and arthropods, whereas exchange among ageing plants declines and communities drift apart. Herbivores arriving late meet plant-specific microbial environments, so microbial mediation of herbivory should be host-specific and seasonally contingent.

## INTRODUCTION

Plant-associated microbial communities shape the health and performance of their hosts, regulating nutrient acquisition, pathogen suppression, defense priming, and tolerance to abiotic stress (Bashir et al., 2022). These communities inhabit the entire plant, from the rhizosphere surrounding the roots to the phyllosphere aboveground (Vorholt, 2012). Yet, phyllosphere communities remain far less understood than their soil counterparts (Wang & Cernava, 2023). The phyllosphere harbors fewer taxa than the rhizosphere, and its lower diversity may indicate reduced functional redundancy and heightened sensitivity to perturbation (Bulgarelli et al., 2013; Lindow & Brandl, 2003; Vorholt, 2012). Its communities also turn over rapidly and exhibit dynamic patterns of community assembly and turnover (Maignien et al., 2014; Morella et al., 2020). Bacteria dominate the phyllosphere (Thapa & Prasanna, 2018), where they colonize a harsh, exposed surface subjected to UV radiation, desiccation, and temperature extremes (Agler et al., 2016; Vorholt, 2012). Far from a passive backdrop, the leaf actively structures the communities it hosts. Surface architecture, including stomatal depressions, trichome bases, and surface topography, creates moist microsites that shelter colonists, while leaf size and shape influence the microclimate those colonists experience (Boyle et al., 2024; Schlechter et al., 2019). Plant chemistry imposes a second filter, as cuticular monoterpenes, flavonoids, and alkaloids facilitate or inhibit bacterial taxa (Bashir et al., 2022; Vorholt, 2012), and secretions at trichome bases can enrich growth-promoting groups (Doan et al., 2020). Because these physical and chemical traits shift with plant ontogeny and across the growing season, the filters themselves are dynamic, potentially restructuring phyllosphere communities through time (Gao et al., 2023; Wang & Cernava, 2023; Xu et al., 2022). Understanding what governs phyllosphere assembly is therefore essential for predicting. Understanding what governs phyllosphere assembly therefore matters beyond the microbes themselves: with limited functional redundancy, changes in community membership may translate directly into changes in the pathogen suppression and defense priming these communities provide.

Community assembly reflects the interplay of deterministic and stochastic processes (Dini-Andreote et al., 2015). Deterministic processes act through selection, as environmental and host filters favor some taxa, whereas stochastic processes include dispersal, ecological drift, and historical contingencies associated with the timing and order of colonization, including priority effects. Null modeling frameworks that compare observed phylogenetic turnover against random expectations can infer the relative contributions of different ecological processes to community turnover (Dini-Andreote et al., 2015; Jia et al., 2022). In the phyllosphere, plant secondary chemistry is a prime candidate for environmental filtering, potentially selecting among taxa across host trait gradients. Yet few habitats are as open to stochastic colonization as the leaf surface. Microbes disperse via wind, rain, soil, and the bodies of insects (Bashir et al., 2022), and this passive, context-dependent immigration can send communities down divergent trajectories. Herbivores may further influence these assembly dynamics by serving as microbial dispersal vectors and by modifying leaf environments through feeding. Their visitation can increase bacterial diversity and reshape community composition (Humphrey et al., 2014; Humphrey & Whiteman, 2020; Ushio et al., 2015), and as their identities and abundances fluctuate, so may the pool of microbial immigrants they carry (dos Santos et al., 2024). Consistent with this openness to repeated immigration, stochastic processes, including ecological drift and homogenizing dispersal, can contribute substantially to phyllosphere community assembly. Despite the growing use of these frameworks across microbial systems, they have rarely been applied to phyllosphere communities in which host chemistry and arthropod composition are both well characterized (Dini-Andreote et al., 2015; Jia et al., 2022).

Milkweeds (genus *Asclepias*) offer an excellent system for testing how phyllosphere communities assemble because host species vary widely in the chemical and physical traits that can filter microbial colonists. Each species carries a characteristic profile of cardenolides, toxic steroidal metabolites present in milkweed latex, ranging from the cardenolide-rich leaves of *A. curassavica* to the nearly cardenolide-free *A. tuberosa* (Agrawal et al., 2012). Although cardenolides are studied primarily as defenses against herbivores, individual compounds also display antimicrobial activity (Agrawal et al., 2012; Akhtar et al., 1992; Huq et al., 1999), making milkweed chemistry a plausible filter on leaf-surface colonists. Consistent with this, phyllosphere bacterial communities differ in composition between *A. curassavica* and *A. syriaca* without a corresponding difference in richness (Hansen & Enders, 2022), a pattern suggesting that cardenolides may reshape which taxa establish rather than how many a leaf supports. Whether such effects hold across a broader cardenolide gradient, and how they interact with seasonal turnover and arthropod-mediated dispersal, remains untested. Species also differ in physical defenses such as trichome density and in the arthropod communities that visit them, which may provide a common and dynamic source of microbial dispersal. This combination allows us to ask how deterministic and stochastic processes jointly shape leaf microbiome assembly across contrasting host species and through time.

Here, we characterized the phyllosphere bacterial communities of four milkweed species, *A. curassavica*, *A. incarnata*, *A. syriaca*, and *A. tuberosa*, which differ in leaf morphology, plant architecture, cardenolide chemistry, trichome density, and associated arthropods, tracking bacterial and arthropod communities across a single growing season. We evaluated the contributions of host identity, plant chemistry, arthropod communities, and time to bacterial diversity and composition, and quantified the relative roles of deterministic and stochastic processes in community assembly. We expected these contributions to differ among hosts and to shift as the season advanced and tested three predictions. First, because cardenolides and trichomes can act as chemical and physical filters on leaf colonists, we predicted that better-defended species (*A. curassavica*, *A. syriaca*) would support less diverse communities and a stronger signature of selection, whereas the nearly cardenolide-free *A. tuberosa* would harbor the most diverse communities, assembled largely through stochastic processes. Second, we predicted a shift from stochastic to deterministic assembly as the season progressed: newly transplanted plants bearing young, sparsely colonized leaves should recruit from a shared environmental pool, producing homogenizing dispersal early on, whereas established communities on ageing leaves should increasingly reflect host-specific selection and, as exchange among plants declines, dispersal limitation. Because chemical filtering differs in strength among hosts, we expected this transition to appear earliest on *A. curassavica* and *A. syriaca* and latest on *A. tuberosa*. Third, because arthropods vector microbes among plants and modify leaf surfaces, we predicted that species hosting more abundant and diverse arthropods, particularly *A. curassavica* and *A. incarnata*, would support higher bacterial diversity and a stronger dispersal signal, rendering arthropod effects host dependent.

## METHODS

### Milkweed species

We studied four milkweed species that span a range of ecological and morphological traits: *Asclepias curassavica*, *A. syriaca*, *A. incarnata*, and *A. tuberosa*. *Asclepias curassavica* (tropical milkweed) is native to the American tropics, spanning the Caribbean, Central America, and northern South America, and is distinguished by its vibrant red-orange flowers and vigorous growth habit (Woodson, 1954). *Asclepias syriaca* (common milkweed) is native to the eastern and central United States, tolerates a wide range of soils and moisture, and bears clusters of pink to purple flowers; as a primary larval host for the monarch butterfly (*Danaus plexippus*), it is central to monarch conservation (DeLaMater et al., 2021). *Asclepias incarnata* (swamp milkweed) occupies wetland habitats such as marshes, floodplains, and riverbanks across the eastern and central United States, where it produces deep pink to purple flowers and abundant nectar for monarchs and other pollinators (Nichter & Gregory, 2018). *Asclepias tuberosa* (butterfly milkweed) grows in prairies, dry meadows, and open woodlands of the central and eastern United States, where a deep taproot confers drought tolerance and bright orange flowers provision adult monarchs during reproduction (Agrawal, 2017).

The four species differ markedly in the chemical and physical defenses most relevant to leaf colonization, spanning a broad range of foliar cardenolides and latex as well as contrasting trichome densities and leaf forms(Agrawal & Fishbein, 2006). These gradients allow us to test the roles of plant chemistry and morphology in phyllosphere assembly.

### Study system and experimental design

We selected four milkweed species in this study: three native species, *A. incarnata*, *A. syriaca* and *A. tuberosa* (sourced from Roundstone Native Seed Company, Upton, KY, USA), and one non-native species: *A. curassavica* (OutsidePride, Salem, OR, USA). We cold-stratified seeds for 14 days, then sowed them into 10-cm diameter pots and grew them in a controlled growth chamber (22°C, 16:8 h light:dark cycle, 55% RH). At three weeks, we fertilized all seedlings with Osmocote slow-release fertilizer (Scotts Company LLC, Marysville, OH, USA). After eight weeks, we swabbed all plants for microbial sampling and transferred them to the University of Alabama Arboretum (Tuscaloosa, AL, USA) on June 11^th^, 2021.

We planted the milkweeds in an old-field area following a randomized block design. Each of 16 blocks received one individual of *A. curassavica*, *A. syriaca*, and *A. tuberosa*, with 50 cm between plants within a block and at least 1 m between blocks. Because of low germination, *A. incarnata* was only included in 8 of the 16 blocks. The design was therefore complete for three species (*n* = 16 each) and incomplete for *A. incarnata* (*n* = 8), giving 56 plants in total. Blocks containing three and four species were interspersed across the field, and block was fitted as a random effect in all models to absorb variation associated with block composition and position. At transplant, we applied a second round of Osmocote and installed drip irrigation, watering for two hours each night except after rainfall.

### Arthropod surveys and plant performance

We conducted weekly surveys from June 17th to August 27th 2021, for a total of 11 surveys, recording all visible arthropods on each milkweed plant. When field identification was not possible, we collected specimens and preserved them in 70% ethanol for later identification in the laboratory. We classified arthropods into feeding guilds: herbivores, predators, omnivores, detritivores, and pollinators. Pollinators were recorded but not analysed further, because few plants flowered (see below). We additionally scored milkweed specialists, taxa that feed principally or exclusively on *Asclepias*, separately from generalist herbivores, since their presence indicates an obligate association with the host rather than incidental occurrence. Counts of the two most frequently encountered specialists, the oleander aphid (*Aphis nerii*) and larvae of the monarch butterfly (*Danaus plexippus*), were recorded separately by species, scored as present or absent on each plant, and analysed individually in addition to their guild totals. Because surveys were visual, arthropods were recorded as present on a plant regardless of whether feeding was observed. Each week, we also measured plant height and counted the number of leaves on every plant to monitor vegetative growth. At the end of the experiment, on September 1st, we harvested aboveground biomass from each plant, dried the samples for four days at 40°C in a walk-in drying oven, and recorded the dry mass. Only 6 of 56 plants flowered during the experiment, all of them *A. tuberosa* or *A. curassavica*, so flowering was not included as a ppredictor,and we used aboveground biomass as a proxy for plant performance rather than reproductive metrics. We did not quantify leaf damage; arthropod abundance served as our measure of herbivore pressure.

### Trichome density and cardenolide content

At the end of the experiment, we collected leaves from each plant for trichome and cardenolide quantification. We quantified trichome density on leaves from all four milkweed species. For each leaf, we counted the number of trichomes on 3 mm^2^ on the abaxial surface under a stereomicroscope (Kariyat et al., 2012). For cardenolides, we used on average 450mg of fully expanded leaf tissue per plant, which we placed in a glassine envelope immediately after collection and dried at 40°C to constant mass. We grounded the dried leaf material to a fine powder and extracted cardenolides in methanol, following protocols established for *Asclepias* species (Cibotti et al., 2025). Samples were analyzed by HPLC using a Zorbax StableBond C18 reversed phase column (5 μm, 150 × 4.6 mm, Agilent Technologies, Santa Clara, CA, USA) and an Agilent 1100 series instrument with diode array detection. The 15 uL injection was eluted at a constant flow of 0.7 mL/min with a gradient of acetonitrile and water as follows: 0–2 min 16% acetonitrile; 25 min 70% acetonitrile; 30 min 95% acetonitrile with a final 8 min hold. Peaks were detected by a diode array detector at 218 nm, and absorbance spectra were recorded from 200–400 nm. Peaks showing a characteristic symmetrical absorption band with a maximum between 217–222 nm were recorded as cardenolides. Sample concentrations were quantified by relating abundances to the peak area of the internal standard (20 μg of digitoxin (Sigma, St. Louis, MO, USA) as internal standard).

### Microbial sample collection, DNA extraction, sequencing, and bioinformatics

To investigate how host plant species influence phyllosphere microbial communities, we sampled leaf surfaces using sterile cotton swabs (MW113; Medical Wire & Equipment Co. Ltd., Corsham, Wiltshire, UK). We swabbed all plants once a month for four months, passing the swab over the surface of each leaf 30 times to ensure consistent sampling. After swabbing, we placed each swab in a 2 mL screw-cap tube and stored it at -80°C until processing. We included two controls that were processed and sequenced alongside the samples: a blank swab as a negative control, and a swab of a gloved hand to account for potential contamination introduced during handling. We extracted DNA using the Qiagen DNeasy Blood & Tissue kit, modifying the manufacturer’s protocol to increase DNA yield following Kueneman et al. (2014). We then PCR-amplified the V4 region of the bacterial 16S rRNA gene using dual-indexed 515F and 806R barcoded primers (Caporaso et al., 2011; Kozich et al., 2013). We purified pooled amplicons with a QIAquick Gel Extraction Kit (Qiagen) and submitted them for sequencing on an Illumina MiSeq at the Tufts University Genomics Core Facility (Boston, MA, USA). The facility provided demultiplexed reads by sample. We imported the forward reads into QIIME2 (Bolyen et al., 2019) to characterize bacterial diversity. We trimmed sequences to 150 bp, quality-filtered them, and denoised them with Deblur (Amir et al., 2017), then clustered the resulting sequences into operational taxonomic units (OTUs) at 97% sequence similarity. We assigned taxonomy to OTUs using a naive Bayes classifier (classify-sklearn) trained on the SILVA reference database and removed chloroplast and mitochondrial sequences along with low-abundance OTUs (those representing <0.005% of total reads) following (Bokulich et al., 2013) We rarefied the OTU table to 3,500 reads per sample, a depth chosen from the rarefaction curves. Sample sizes varied across the four sampling months (Table S1).

### Processes governing phyllosphere community assembly across the season

To determine the community assembly processes structuring bacterial composition, we used a null modeling framework based on phylogenetic and abundance information following Ecological succession and stochastic variation in the assembly of *Arabidopsis thaliana* phyllosphere communities (Dini-Andreote et al., 2015; Stegen et al., 2013; Stegen et al., 2015). In brief, phylogenetic turnover between communities was quantified as the β mean nearest taxon distance (βMNTD) across all samples, calculated with the function “comdistnt” (abundance.weighted = TRUE) in the R package *picante* (Kembel et al., 2010). Null distributions of βMNTD were generated by randomly shuffling taxa across the tips of the phylogenetic tree (999 permutations), thereby removing the signal of deterministic ecological selection. The β-nearest taxon index (βNTI) was then calculated as the standardized deviation between observed and null βMNTD values. Values of βNTI > +2 and < −2 indicate significant deviations from the null expectation, reflecting the dominance of variable and homogeneous selection, respectively, whereas values between −2 and +2 indicate the dominance of stochastic assembly processes. To further evaluate these stochastic processes, we calculated the Raup-Crick metric based on Bray-Curtis dissimilarity (RCbray) using 999 permutations. RCbray values > +0.95 indicate dispersal limitation coupled with drift, whereas values < −0.95 indicate that community turnover is driven by homogenizing dispersal. Finally, intermediate values (−0.95 < RCbray < +0.95) were interpreted as undominated processes. These categories describe what drove turnover between a pair of communities, not the state of a single community. Homogeneous selection means a consistent filter, such as shared host chemistry, favored the same lineages on both plants, making them more similar than chance. Variable selection means contrasting conditions favored different lineages, driving the two apart. Homogenizing dispersal means organisms moved between plants often enough to override local differences. Dispersal limitation, formally dispersal limitation coupled with drift, means few organisms moved between plants, so each community changed independently through colonization and local extinction and the two diverged. Undominated turnover means none of these crossed the detection thresholds; it does not indicate drift specifically, but encompasses weak selection, moderate dispersal, drift, and diversification acting together. Selection is deterministic, whereas dispersal and drift are stochastic with respect to taxon identity. The relative contribution of each process was quantified as the proportion of pairwise comparisons assigned to each category. For full details of this method, see Stegen et al. (2013) and Dini-Andreote et al. (2015).

### Statistical analyses

All analyses were conducted in R v. 4.6.0 (R Core Team, 2026) using generalized linear mixed models (GLMMs) fitted by maximum likelihood in *glmmTMB* v. 1.1.14 (Brooks et al., 2017), unless stated otherwise. All models included host plant species (*A. tuberosa*, *A. incarnata*, *A. syriaca*, and *A. curassavica*) as a fixed effect, with model-specific random effect structures noted below and summarized in Table S2. Error distributions were chosen by data type and confirmed with simulation-based residual diagnostics in *DHARMa* v. 0.4.7 (Hartig, 2022), testing residual uniformity (Kolmogorov-Smirnov), dispersion, zero-inflation, and outliers. For count responses (richness, abundance), we used Poisson or, when overdispersed, negative binomial distributions, whereas for continuous responses (Shannon diversity, Pielou’s evenness, Faith’s phylogenetic diversity, and cardenolide concentration), we used Gaussian (with log or log(x + 1) transformation), lognormal, or Tweedie distributions, whichever satisfied the diagnostics (Table S2). Fixed effects were tested with type III Wald χ² tests *car* (Fox & Weisberg, 2019), with sum-to-zero contrasts applied to models containing interactions. Significant effects were followed by Tukey or Sidak-adjusted pairwise comparisons of estimated marginal means (*emmeans* v. 2.0.3 (Lenth & Piaskowski, 2026), computed within each month for seasonal analyses and back-transformed for presentation. Compact letter displays (α = 0.05) were generated with *multcompView*.

### Cumulative models

Arthropod richness, abundance, Shannon diversity (H′), and Pielou’s evenness (J′) were calculated from the season-summed community of each plant (per-taxon counts pooled across all 11 surveys) and modelled with host species as a fixed effect and experimental block as a random effect, using the distributions stated above. Differences in arthropod community composition among host species were tested on this same cumulative dataset with permutational multivariate analysis of variance (PERMANOVA; *adonis2*, *vegan* v. 2.3.7, Oksanen (2008); 999 permutations), computed on Jaccard dissimilarities of presence/absence data because the cumulative counts were dominated by a few taxa and we focused on taxon turnover. Pairwise differences among species were assessed with pairwise PERMANOVA, and community structure was visualized with non-metric multidimensional scaling (NMDS) on the same dissimilarity matrix.

### Temporal models

Arthropod alpha diversity metrics (richness, abundance, Shannon diversity, and evenness) were additionally modelled across the season, with host species, month, and their interaction (species × month) as fixed effects, and experimental block as a random effect; species were compared within each month using the pairwise procedure described above.

Phyllosphere bacterial alpha diversity (richness, Shannon diversity, Pielou’s evenness, and Faith’s phylogenetic diversity) was modelled from the rarefied community (3,500 reads per sample) across the four sampling months, with host species, month, and their interaction (species × month) as fixed effects, and random effects of plant identity (to account for repeated sampling of the same plant) and experimental block. Shannon diversity and evenness were modelled with a Gaussian distribution, richness with a negative binomial distribution, and Faith’s phylogenetic diversity with a lognormal distribution; responses were log-transformed where diagnostics required it.

### Bacterial community composition

Differences in bacterial community composition among host species were tested with PERMANOVA (*adonis2*, 999 permutations) on Bray-Curtis dissimilarities of the rarefied August community. Pairwise differences among species were assessed with pairwise PERMANOVA, with p-values adjusted by the Bonferroni method, and community structure was visualized with NMDS on the same dissimilarity matrix. A heatmap was generated in *ggplot2* to visualize relative abundances of bacterial taxa across milkweed species, using a color gradient proportional to taxon abundance.

### Effects of arthropods on the bacterial community

To test whether arthropods reshaped the bacterial community, bacterial OTU richness and Shannon diversity in the late-season (August) sampling were modelled as a function of arthropod predictors. Separate GLMMs were fitted with overall arthropod richness, herbivore richness, oleander aphid abundance, and monarch larval abundance, each entered as a fixed effect together with host species and their interaction, and with block as a random effect. OTU richness was modelled using a negative binomial distribution, and Shannon diversity using a lognormal distribution.

### Contribution of arthropods and cardenolides to phyllosphere bacterial community

Within each host species, the association between herbivore presence (monarch larvae and oleander aphids, scored as presence/absence) and bacterial responses (*Methylobacteriaceae* relative abundance and bacterial OTU richness) was evaluated for the August sampling using Spearman rank correlations, which for a binary predictor and a continuous response provide the point-biserial form of the rank correlation.

Variation among host species in foliar cardenolide chemistry was characterized first, to quantify the number of cardenolides each plant produced. We resolved four distinct cardenolide peaks across all plants. The fourth occurred in only two plants and was excluded from analysis, leaving three compounds analysed individually alongside total cardenolide concentration. Total cardenolide concentration and three individual cardenolides were each analyzed with Gaussian GLMMs, with host species as the fixed effect and block as random effect. Total cardenolides and cardenolide 3 were log (x + 1) transformed to meet parametric assumptions, while cardenolides 1 and 2 were modelled untransformed.

### Relationships between plant chemistry, and the bacterial community

To test the effects of hosts on the bacterial community and on arthropods, we fitted GLMMs containing cardenolide concentration, host species, and their interaction as fixed effects. *Asclepias tuberosa* was excluded from these models because it lacked detectable cardenolides, and *A. syriaca* was additionally excluded from the model for cardenolide 3. To assess whether accounting for host relatedness altered the estimated effects of cardenolides, we also included a two-level grouping of the host species into clades following (Fishbein, 2011): the Incarnatae clade (*A. curassavica* and *A. incarnata*) and the Temperate North American clade (*A. syriaca* and *A. tuberosa*). Including clade did not improve model fit or alter the estimated cardenolide effects, so we report the models without it. For the bacterial response (September OTU richness, modelled using a Poisson or negative binomial distribution), separate models were fitted for total cardenolides and for each of the three individual cardenolides. For the arthropod responses, total cardenolides were used to test the effects on arthropod richness (Poisson), oleander aphid abundance (negative binomial), and monarch larval abundance (Poisson). In every case the host-species-specific slopes were estimated and compared with the emtrends function, using adjusted contrasts of slopes.

## RESULTS

### Seasonal dynamics and taxonomic succession in the milkweed phyllosphere

Bacterial richness differed significantly among host plant species (χ² = 21.6, df = 3, p < 0.001, Fig. 1a), across sampling months (χ² = 15.3, df = 2, p < 0.001, Fig. 1a), and with the host species-by-month interaction (χ² = 28.8, df = 6, p < 0.001, Fig. 2a). By September, *A. tuberosa* supported the highest bacterial richness (239 ± 10.2 OTUs, mean ± SE), followed by *A. syriaca* (186 ± 9.9), *A. incarnata* (144 ± 12.3), and *A. curassavica* (140 ± 6.22). Faith’s phylogenetic diversity exhibited similar responses, varying among host plant species (χ² = 23, df = 3, p < 0.001, Fig. S1), across sampling months (χ² = 75.3, df = 3, p < 0.001), and with the host species-by-month interaction (χ² = 42.7, df = 9, p < 0.001). Both bacterial richness and Faith’s phylogenetic diversity increased over the growing season, reaching their highest levels during the final sampling month across all host species (Fig. 1a; Fig. S1).

**Fig. 1.**
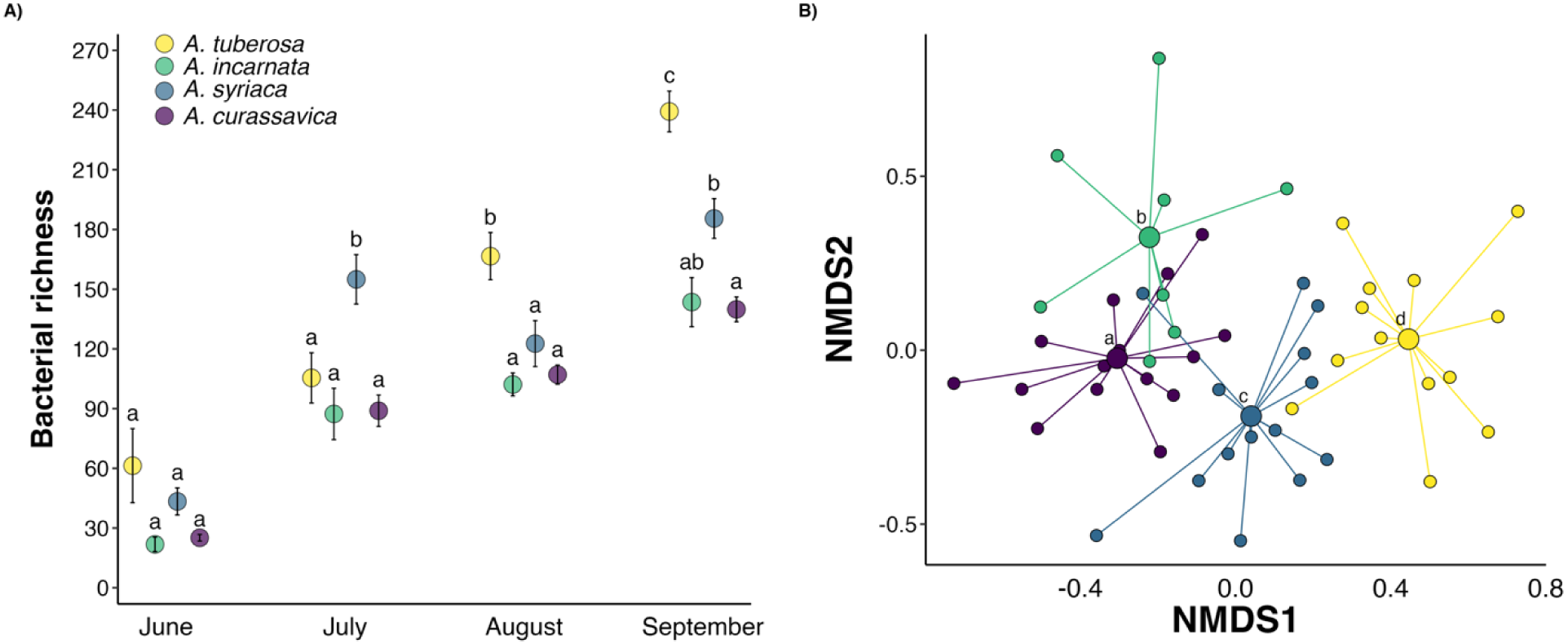
Bacterial community responses to host plant species across the growing season. (A) Bacterial richness (OTUs), and (B) nonmetric multidimensional scaling (NMDS) ordination of bacterial OTU composition during the final sampling month. Error bars represent means ± 1 SE. Different letters indicate significant differences at p < 0.05, determined by post hoc comparisons of estimated marginal means with Sidak adjustment. Different colors represent host plant species (purple = *A. curassavica*, blue = *A. syriaca*, green = *A. incarnata*, and yellow = *A. tuberosa*).

**Fig. 2.**
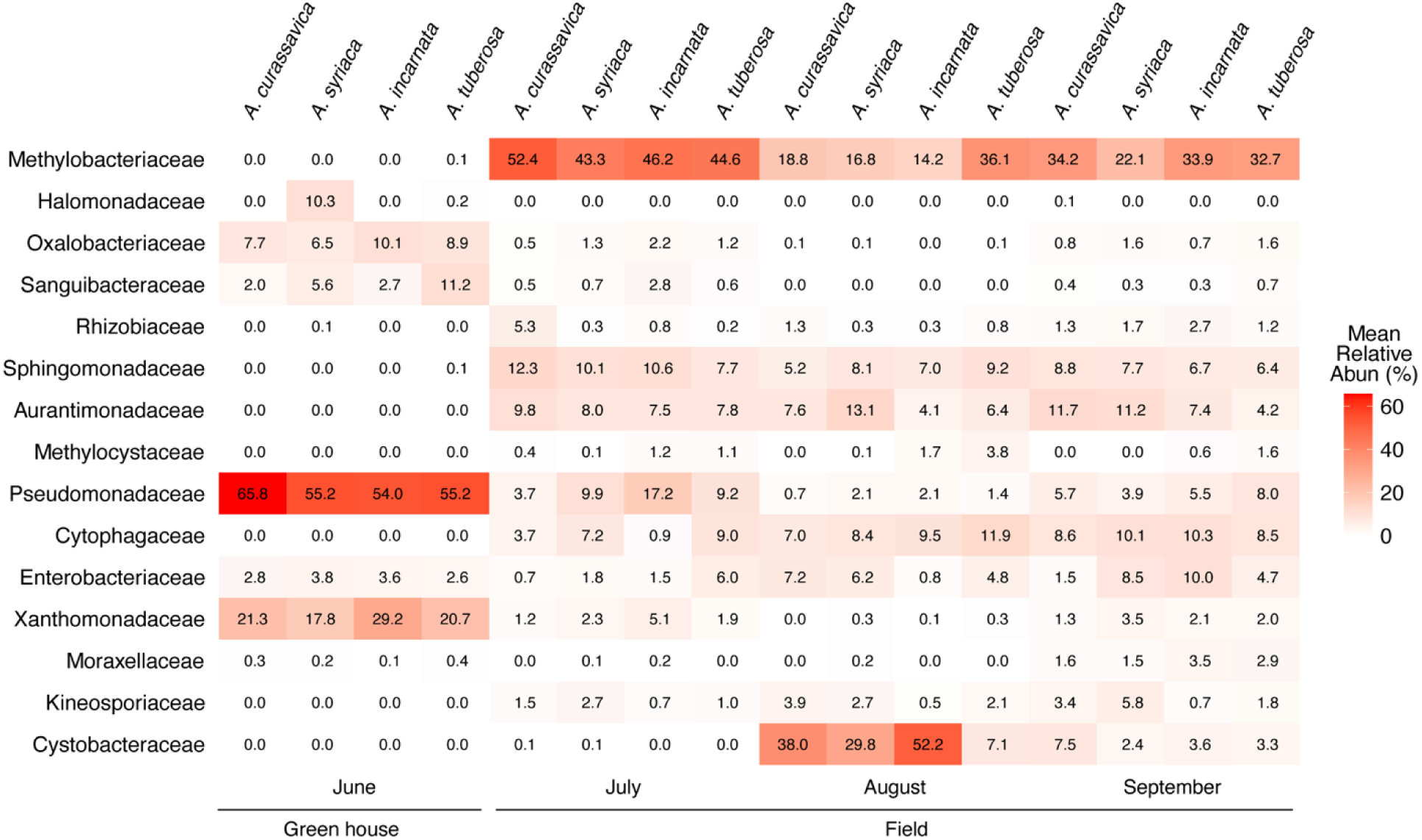
Heatmap illustrating bacterial community composition at the family level across four milkweed species: *Asclepias curassavica*, *A. syriaca*, *A. incarnata*, and *A. tuberosa*. Color intensity reflects the relative abundance of bacterial families, with darker shades indicating higher relative abundance.

Bacterial community composition differed significantly among host species (PERMANOVA: F = 2.9, R² = 0.15, p < 0.001, Fig. 1b). All four host species harbored distinct bacterial communities, except *A. curassavica* and *A. incarnata*, which did not differ significantly (Fig. 2b).

To visualize differences in bacterial community composition among host plant species, we generated a heatmap depicting the relative abundances of bacterial families associated with each milkweed species (Fig. 2). Prior to field transplantation, phyllosphere bacterial communitie were dominated by Pseudomonadaceae and Xanthomonadaceae. Following field transplantation, the relative abundances of these families declined sharply across all species. In contrast, relative abundance of Methylobacteriaceae was 44% higher one month after field transplantation across all host species (Fig. 2).

### Arthropod community

Milkweed species differed in both total arthropod abundance and community composition (Table S3). Per plant, arthropod abundance was highest on *A. incarnata* (1,060 individuals per plant) and lowest on *A. syriaca* (280), and non-aphid abundance was likewise highest on *A. incarnata* (17% per plant) and lowest on *A. tuberosa* (8%). Raw totals are reported in Table S1; note that *A. incarnata* was grown in only 8 blocks.

Cumulative arthropod richness differed among the four milkweed species (χ² = 29.3, df = 3, p < 0.001, Fig. 3). Richness was highest on *A. incarnata* (9.5 ± 1.24 species, mean ± SE), followed by *A. curassavica* (6.88 ± 0.74), *A. tuberosa* (3.94 ± 0.75), and *A. syriaca* (3.88 ± 0.83). Across the season, richness varied with sampling month (χ² = 27.8, df = 3, p < 0.001, Fig. 3b), however, the species × month interaction was not significant, indicating no evidence that differences among host species changed over time (χ² = 14.9, df = 9, p = 0.09). Shannon diversity and Pielou’s evenness were not significantly associated with host species (Shannon: χ² = 2.3, df = 3, p = 0.40; evenness: χ² = 4.4, df = 3, p = 0.20, Fig. S1a, b), month (Shannon: χ² = 1.3, df = 3, p = 0.71; evenness: χ² = 2.1, df = 3, p = 0.54, Fig. S1a, b), or their interaction (Shannon: χ² = 5.2, df = 9, p = 0.80; evenness: χ² = 2.3, df = 9, p = 0.98, Fig. S1a, b). Together, these results indicate that milkweed species differed in arthropod abundance and richness overall, whereas evenness and Shannon diversity remained similar among hosts over the season.

**Fig. 3.**
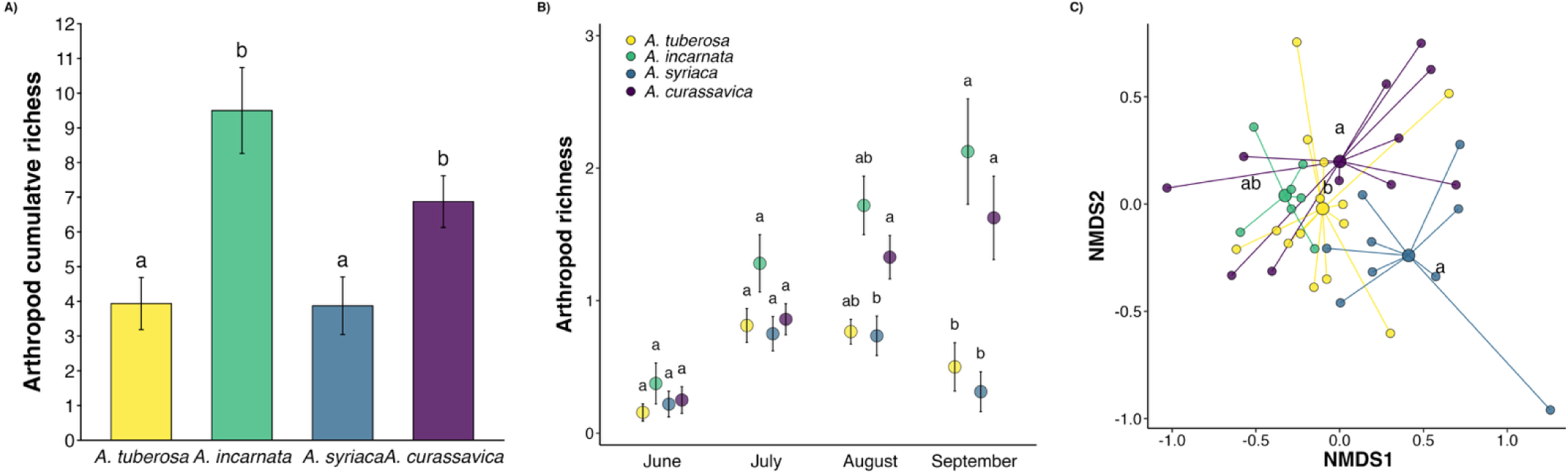
Arthropod richness and community composition across four milkweed species. (a) Cumulative arthropod richness, calculated for each plant as taxon richness across the entire season and presented as the mean across plants. (b) Arthropod richness by month, calculated for each plant as its mean weekly taxon richness within each month. In (A) and (B), error bars show the mean ± 1 SE, and different letters denote significant differences among host species (p < 0.05) based on Sidak-adjusted pairwise comparisons of estimated marginal means; in (B), species are compared within each month. (C) Non-metric multidimensional scaling (NMDS) ordination of arthropod community composition for the cumulative data, based on Jaccard dissimilarities of presence/absence data (stress = 0.19). Each point represents an individual plant, color-coded by host species, with larger symbols indicating group centroids and segments connecting plants to their respective centroids. Different letters denote significant differences in community composition among host species based on pairwise PERMANOVA. Different colors represent host plant species (purple = *A. curassavica*, blue = *A. syriaca*, green = *A. incarnata*, and yellow = *A. tuberosa*).

Arthropod community composition varied significantly among the four milkweed species (PERMANOVA: F = 2.9, R² = 0.19, df = 3, p = 0.001, Fig. 3c). Arthropod communities associated with *A. curassavica* and *A. tuberosa* differed significantly (p = 0.044, Fig. 3c), and *A. syriaca* also differed significantly from *A. tuberosa* (p = 0.042, Fig. 3c). However, *A. tuberosa* and *A. incarnata* did not differ significantly in arthropod community composition (p = 0.30, Fig. 3c).

### Contribution of arthropods and cardenolides to phyllosphere bacterial communities

To evaluate potential mechanisms underlying variation in phyllosphere bacterial richness, we examined its associations with host identity, plant defensive chemistry (cardenolides), and arthropod visitors. The interaction between milkweed species and arthropod richness was significant (χ² = 27.4, df = 3, p < 0.001, Fig. 4a), indicating that the relationship between arthropod richness and bacterial richness differed among milkweed species. Bacterial richness also differed among milkweed species (χ² = 22.2, df = 3, p < 0.001, Fig. 4a), whereas the interaction no significant association between arthropod richness and bacterial richness (χ² = 1.5, df = 1, p = 0.2, Fig. 4a).

**Fig. 4.**
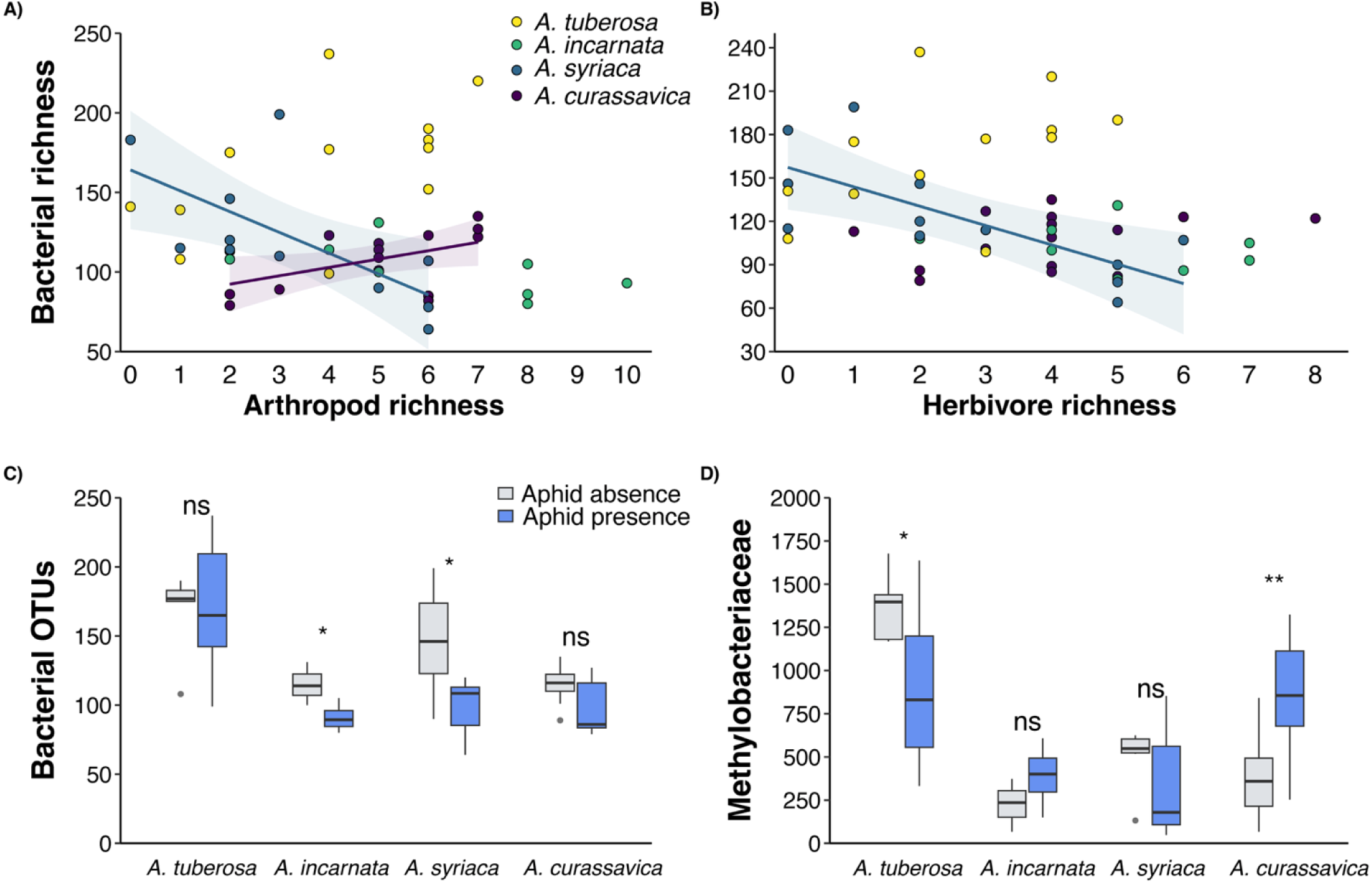
Relationships between phyllosphere bacterial community metrics and arthropod visitors across milkweed species. (A, B) Regression plots showing how (A) arthropod richness and (B) Association between herbivore richness and bacterial OTU richness, with lines and shaded areas representing regressions and 95% confidence intervals. (C, D) Boxplots comparing (C) bacterial OTU richness and (D) *Methylobacteriaceae* relative abundance among plants with (blue) and without (gray) oleander aphids (*Aphis nerii*) present. Boxes indicate the interquartile range (IQR), lines show medians, and whiskers extend to 1.5× IQR. Aphid presence was associated with reduced bacterial richness and altered *Methylobacteriaceae* abundance in some host species.

**Fig. 5.**
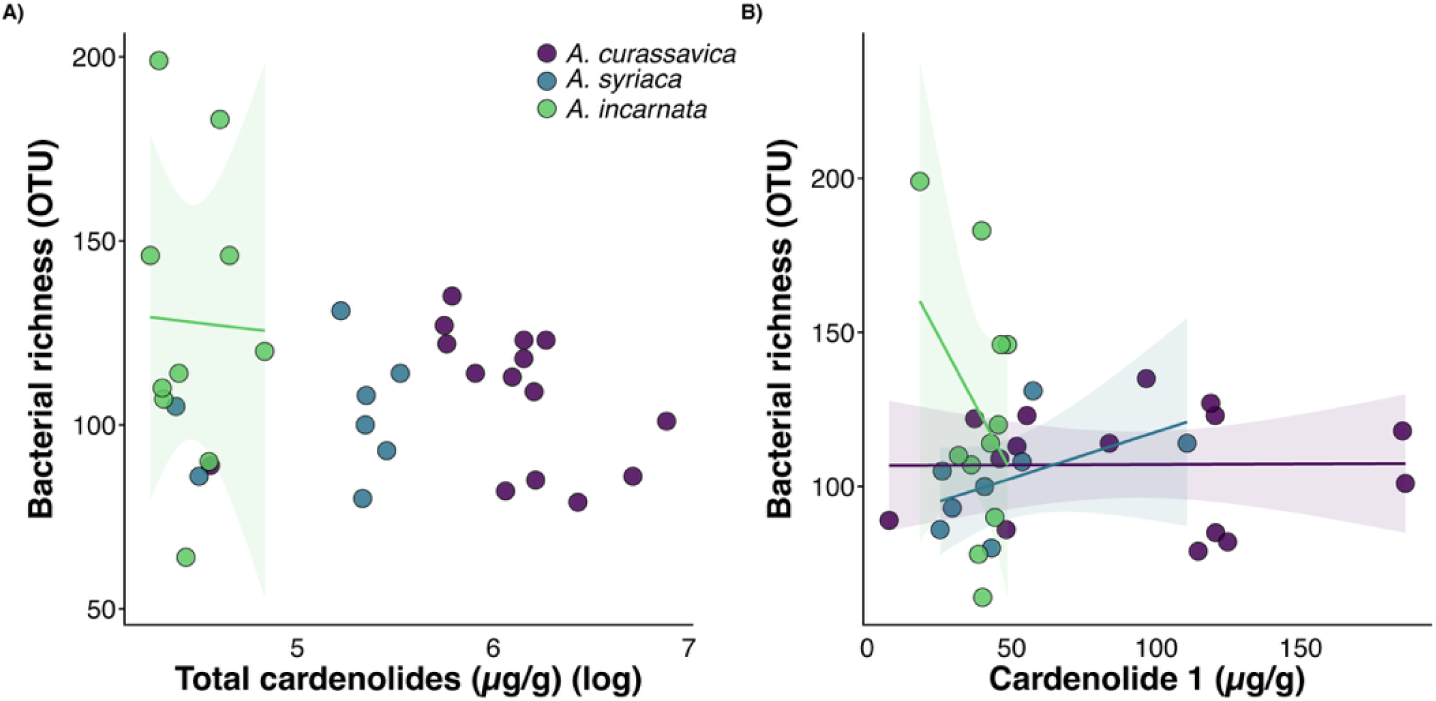
Relationships between cardenolide concentration and bacterial OTU richness among milkweed species. (A) Bacterial richness as a function of total cardenolide concentration (log µg/g) and (B) cardenolide 1 concentration (µg/g) for *A. curassavica* (purple), *A. syriaca* (blue), and *A. incarnata* (green). Points represent individual samples. Lines and shaded areas represent regressions and 95% confidence intervals.

For each arthropod metric, the species-by-arthropod interaction was statistically significant. The species-by-herbivore richness interaction was significant (χ² = 21.1, df = 3, p < 0.001, Fig. 4b), as were the species-by-aphid abundance interaction (χ² = 21.1, df = 3, p < 0.001, Fig. 4c) and the species-by-larval abundance interaction (χ² = 11.3, df = 3, p = 0.009, Fig. 4d). In each model, bacterial richness differed among milkweed species (herbivore: χ² = 14.2, df = 3, p = 0.002; aphid: χ² = 29.5, df = 3, p < 0.001; larvae: χ² = 26.6, df = 3, p < 0.001, Table S4), whereas no significant overall association was detected for the corresponding arthropod metric (herbivore richness: χ² = 1.7, df = 1, p = 0.1, Fig. 4b; aphid abundance: χ² = 0.4, df = 1, p = 0.5, Fig. 4c. Together, these results show that associations between arthropod metrics and phyllosphere bacterial richness were host-specific, with *A. syriaca* in particular showing negative relationships between arthropod metrics and bacterial richness.

The association of cardenolides with bacterial richness differed among milkweed species (χ² = 13.45, df = 2, p = 0.001, Fig. 6a). *A. incarnata* showed a negative association between total cardenolide concentration and OTU richness, whereas no significant associations were detected for *A. syriaca* or *A. curassavica*. For cardenolide 1, the species-by-cardenolide interaction was also statistically significant (χ² = 19.33, df = 2, p < 0.001, Fig. 6b); *A. curassavica* showed a positive association, whereas *A. incarnata* and *A. syriaca* showed negative associations. Cardenolide 2 also exhibited a significant species-dependent interaction, whereas no significant interaction was detected for cardenolide 3 (Fig. S3).

**Fig. 6.**
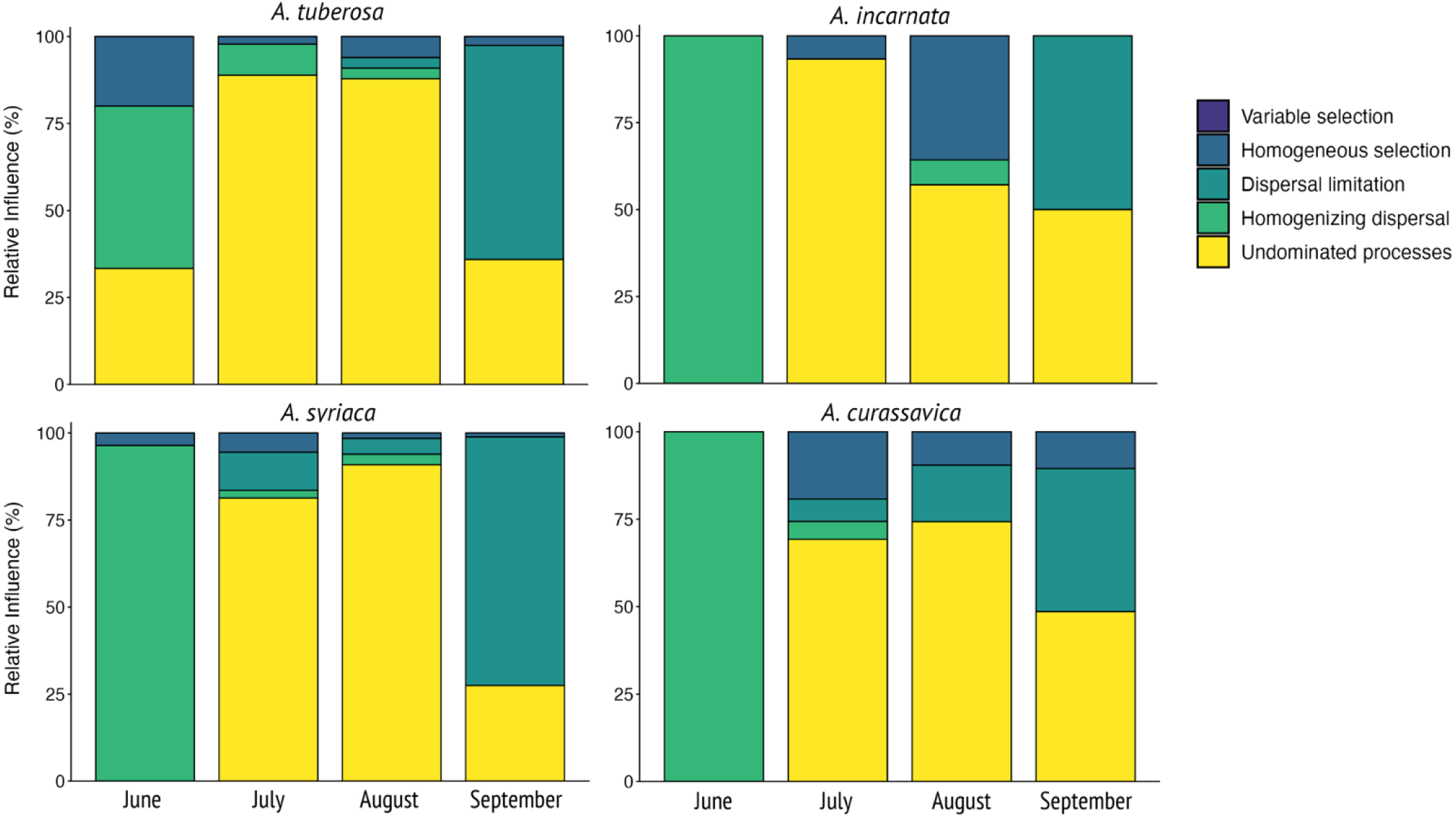
Relative contribution of assembly processes structuring the phyllosphere bacterial communities among four milkweed species. Bar plots show the relative contributions of assembly processes to community turnover at four sampling times (i.e., June, July, August, and September). βNTI and RCBray values were used to infer the relative contribution of each assembly process. Colors represent the inferred assembly processes, i.e., variable selection, homogeneous selection, dispersal limitation, homogenizing dispersal, and undominated processes. Variable selection is shown for completeness but was not detected in any species or month.

### Processes governing phyllosphere community assembly across the season

The relative contributions of stochastic and deterministic processes to bacterial community assembly shifted across the growing season, with patterns varying among milkweed species (Fig. 6). In June, homogenizing dispersal was the dominant assembly process for *A. syriaca* (96.4%), *A. incarnata* (100%), and *A. curassavica* (100%). In *A. tuberosa*, June communities were more evenly divided among homogenizing dispersal (46.7%), undominated processes (33.3%), and homogeneous selection (20.0%). Throughout July and August, undominated processes prevailed across all four species, ranging from 57.1 to 93.3% of pairwise community comparisons (*A. syriaca*: 81.3 to 90.9%; *A. tuberosa*: 87.9 to 88.9%; *A. curassavica*: 69.2 to 74.3%; *A. incarnata*: 57.1 to 93.3%), indicating that, for most pairwise comparisons, no single assembly process crossed the thresholds used to infer selection, homogenizing dispersal, or dispersal limitation. By September, a marked shift toward dispersal limitation was observed in all species, strongest in *A. syriaca* (71%) and *A. tuberosa* (62%), and more moderate in *A. incarnata* (50%) and *A. curassavica* (41%). Homogeneous selection contributed modestly and inconsistently across the season, reaching its highest values in *A. tuberosa* in June (20%), *A. curassavica* in July (19%), and *A. incarnata* in August (36%). The value observed for *A. incarnata* in August was the largest contribution of homogeneous selection observed for any species or month, whereas this process remained below 6% in *A. syriaca* throughout the season. Variable selection did not contribute to community turnover at any time point for any species.

## Discussion

Understanding how microbial communities assemble on plant surfaces remains a central challenge in microbial ecology. Yet, few studies have simultaneously examined the interplay of deterministic and stochastic assembly processes across multiple host species and through time. By monitoring phyllosphere bacteria, and arthropod communities across four *Asclepias* species over a growing season, we found that bacterial community assembly was jointly structured by host identity and seasonally dynamic ecological processes. Host identity consistently structured bacterial communities, inferred assembly processes shifted from early homogenizing dispersal toward late-season dispersal limitation, and arthropods contributed additional host-specific variation. Together, these findings suggest that phyllosphere microbiomes are assembled through dynamic interactions between host traits, temporal changes in dispersal and selection, and arthropod associations, shedding new light on the assembly of plant-associated microbial communities.

### Host identity as a primary determinant of phyllosphere bacterial community structure

The differences in bacterial richness, phylogenetic diversity, and community composition among the four *Asclepias* species are consistent with the well-established role of host plant identity as an important determinant of phyllosphere community assembly (Bulgarelli et al., 2013; Lajoie & Kembel, 2021; Vorholt, 2012). By the end of the growing season, *A. tuberosa* supported the highest bacterial richness, followed by *A. syriaca*, *A. incarnata*, and *A. curassavica*. Although A. tuberosa has been reported to contain low foliar cardenolide concentrations (0.52 mg/g; Agrawal & Fishbein, 2006), we detected none in our plants, consistent with the low concentrations typical of this species under our growing conditions. This ranking broadly corresponded to differences in cardenolide chemistry: *A. tuberosa*, with no detectable cardenolides in our study, harbored the richest bacterial community, whereas *A. curassavica*, characterized by comparatively high cardenolide concentrations, supported lower richness. This pattern is consistent with the possibility that plant secondary metabolites contribute to filtering microbial colonists, as reported in other systems (Schlechter et al., 2019; Thoenen et al., 2023). However, because host species differ simultaneously in multiple chemical, morphological, and phenological traits, cross-species patterns alone cannot isolate cardenolides as the causal mechanism underlying differences in bacterial richness.

Secondary metabolites, including terpenoids and alkaloids, can suppress or enrich distinct bacterial taxa through both direct antimicrobial activity and indirect modulation of host immune signaling (Jacoby et al., 2021; Vorholt, 2012). Cardenolides are steroidal glycosides known to inhibit Na^+^/K^+^-ATPase activity broadly across eukaryotes, but their antimicrobial potential against prokaryotes remains largely unexplored. In our study, associations between cardenolide concentration and bacterial OTU richness were strongly species-dependent. In *A. incarnata,* increasing total cardenolide concentration was associated with reduced bacterial richness, whereas individual cardenolide compounds exhibited contrasting relationships with bacterial richness across host species. These results suggest that if cardenolides contribute to phyllosphere bacterial community structure, their effects may depend on both compound identity and host context rather than total concentration alone.

### Seasonal dynamics and taxonomic succession in the milkweed phyllosphere

Bacterial richness and phylogenetic diversity increased consistently over the growing season across all four milkweed host species, reaching their highest values at the end of the experiment. This temporal pattern is consistent with phyllosphere communities assembling through a cumulative, successional process driven by repeated immigration events from environmental reservoirs, including air, soil, rain, and visiting organisms (Maignien et al., 2014; Morella et al., 2020; Vorholt, 2012). As leaves age, changes in surface chemistry, cuticle integrity, and metabolite leaching may alter resource availability and microsite heterogeneity, potentially allowing a broader range of microbial taxa to establish over time (Gao et al., 2023; Wang & Cernava, 2023). This late-season rise in richness coincides with the shift toward dispersal limitation in our assembly analyses, indicating that increasing local diversity occurred alongside greater differentiation in the inferred processes governing community turnover. Although dispersal limitation can promote divergence among local communities, our data do not establish that it caused the observed increase in richness; both patterns may instead reflect concurrent changes in plant age, environmental conditions, source pools, or colonization history. Prior to field exposure, the phyllosphere communities were dominated by Pseudomonadaceae and Xanthomonadaceae, two widespread families of plant-associated bacteria (Morella et al., 2020). Within one month of field exposure, the relative abundance of these families declined by approximately 53 to 65% across all species, while Methylobacteriaceae increased markedly, rising by approximately 44% across all plant species. This succession is consistent with rapid restructuring of phyllosphere communities following exposure to field conditions and broader environmental source pools (Leducq et al., 2022; Morella et al., 2020). Methylobacteriaceae are among the most prevalent bacterial groups in the phyllosphere across diverse plant hosts. As facultative methylotrophs, they can use methanol released during cell wall pectin demethylation as a carbon source, and some members have been associated with plant growth promotion, pathogen suppression, and stress tolerance (Glick, 2014; Leducq et al., 2022; Spaepen et al., 2007; Vorholt, 2012). Their increase after field transplant may therefore reflect changes in plant physiology, environmental exposure, or both, although the underlying mechanism was not directly tested here. Because temporal shifts in Methylobacteriaceae varied among host species and were associated with aphid presence in a host-dependent manner, this family illustrates how responses of dominant phyllosphere lineages can depend on interacting host and biotic contexts. Within this family, Leducq et al. (2022) documented a seasonal turnover in temperate forests from high-yield strategists, which convert substrate efficiently into biomass, toward rapid-growth strategists, which prioritize fast division, further indicating that succession operates even at the within-family level as plant physiology shifts.

### Processes governing phyllosphere community assembly across the growing season

A central contribution of this study is the quantification of assembly processes structuring phyllosphere bacterial communities across the growing season (Stegen et al., 2013; Stegen et al., 2015). Our results reveal a clear seasonal trajectory in the relative contributions of inferred assembly processes (Dini-Andreote et al., 2015; Jia et al., 2022). Early in the growing season, following transplant, homogenizing dispersal dominated assembly in three of the four species (*A. syriaca*, *A. incarnata*, and *A. curassavica*), consistent with high rates of microbial exchange or exposure to shared source pools reducing compositional differentiation among plants (Bechtold et al., 2024; Maignien et al., 2014). The common greenhouse history of these plants before field transplantation may also have contributed to this initial similarity, making it difficult to disentangle the effects of shared source pools from those of early plant developmental stage.

Throughout July and August, undominated processes prevailed across all species (57 to 93% of assembly), indicating that pairwise community turnover could not be attributed to strong selection, homogenizing dispersal, or dispersal limitation under the thresholds used in the null-model framework. Importantly, this category should not be interpreted as direct evidence that ecological drift alone governed community composition; rather, it may encompass weak selection, moderate dispersal, drift, diversification, or combinations of processes that do not generate sufficiently strong deviations from null expectations. The predominance of undominated processes therefore suggests a mid-season transition in which no single inferred process consistently dominated community turnover. By the end of the season, all species shifted toward dispersal limitation, most strongly in *A. syriaca* (71%) and *A. tuberosa* (62%), and more moderately in *A. incarnata* (50%) and *A. curassavica* (41%). This pattern is consistent with increasing spatial differentiation among local communities as the season progressed. Several non-exclusive mechanisms could contribute to this transition, including changes in abiotic or arthropod-mediated dispersal, divergence in leaf traits and chemistry, shifts in environmental source pools, and the accumulation of distinct colonization histories. Distinguishing among these mechanisms will require direct measurements of microbial dispersal and source pools (Smets et al., 2023). Variable selection did not contribute to community turnover at any time point in any species, providing little evidence that heterogeneous environmental conditions generated strong phylogenetically detectable divergent selection within host species at the temporal scales examined. Where deterministic selection was detected, it took the form of homogeneous selection, and even this was limited and inconsistent: low throughout the season in *A. syriaca* (never exceeding 6%), but appearing as temporally restricted increases in the other three species, i.e., in *A. tuberosa* in June (20%), *A. curassavica* in July (19%), and most strongly in *A. incarnata* in August (36%), the largest such signal observed in this study. Rather than a gradual, season-long increase in selective pressure, these results suggest that phylogenetically detectable homogeneous selection was episodic and host-specific rather than increasing consistently as the season progressed.

The species-specific trajectories further show that no single host trait readily explained the timing or magnitude of these transitions. *A. syriaca* and *A. tuberosa* exhibited the strongest late-season dispersal limitation despite contrasting in cardenolide concentration and other leaf traits, whereas *A. curassavica*, characterized by comparatively high cardenolide concentrations, showed a more moderate late-season shift. Similarly, the largest contribution of homogeneous selection occurred in *A. incarnata* in August rather than in the host with the highest cardenolide concentration. These patterns reinforce the conclusion that cardenolide concentration alone does not predict the inferred balance of selection and dispersal. Instead, multiple host attributes, including chemistry, morphology, phenology, and associated arthropod communities, may jointly contribute to species-specific assembly trajectories.

### Context-dependent effects of arthropods on phyllosphere bacterial communities

Our results reveal that associations between arthropod metrics and phyllosphere bacterial richness were strongly dependent on host plant identity. These findings are consistent with the growing body of evidence that arthropod visitation represents a biologically meaningful source of phyllosphere microbiome variation (dos Santos et al., 2024; Humphrey et al., 2014; Humphrey & Whiteman, 2020; Ushio et al., 2015). The significant species-by-arthropod interactions indicate that relationships between arthropod visitation and bacterial diversity cannot be generalized across host species. Host-specific differences in chemistry, morphology, or plant responses to herbivory may contribute to these contrasting relationships, although the present observational design does not isolate their respective roles. Aphid associations provide the clearest example of this host dependence. Aphids can influence phyllosphere microbial communities through multiple non-exclusive pathways. Honeydew deposition, for example, provides a concentrated, sugar-rich substrate that can stimulate microbial growth (Stadler & Müller, 1996; Wolfgang et al., 2023), feeding wounds may create entry points for environmental bacteria, and aphid-induced plant signaling responses, including pathways involving jasmonic acid and salicylic acid, can alter the chemical composition of the leaf surface in ways that can differentially favor or exclude specific microbial taxa (Glazebrook, 2005; Humphrey et al., 2014). The positive association between aphid presence and bacterial Shannon diversity in *A. curassavica* and *A. tuberosa* in July is consistent with the possibility that honeydew deposition increases resource availability to epiphytic bacteria. In contrast, the negative association between aphids and *A. syriaca* in August could arise from contrasting changes in resource availability, competitive interactions, plant physiology, or defense signaling.

Aphid associations on Methylobacteriaceae illustrate the complexity of these interactions. As facultative methylotrophs that can use plant-derived methanol (Leducq et al., 2022; Vorholt, 2012), these bacteria may respond to changes in plant physiology. At the same time, honeydew deposition may alter resource competition by favoring bacteria capable of exploiting readily available sugars. The contrasting associations observed among host species therefore point to context dependence rather than a uniform aphid effect. This interpretation is consistent with evidence that insect herbivory can restructure native leaf microbiomes in species- and genotype-dependent ways (Humphrey & Whiteman, 2020). Monarch larval abundance did not significantly affect bacterial richness, providing no evidence for a detectable relationship under the conditions observed here. However, monarch densities were low throughout the study, potentially limiting statistical power and the magnitude of herbivory-induced changes. Thus, the absence of a significant association should not be interpreted as evidence that monarch herbivory cannot influence phyllosphere communities under greater herbivore pressure.

Our arthropod results should be interpreted in light of the overwhelming numerical dominance of *Aphis nerii*, which accounted for over 98% of all individuals recorded. Non-aphid arthropods totalled 651 individuals across the season, so our measures of arthropod richness and abundance largely reflect the ecology of a single specialist herbivore rather than the milkweed arthropod community as a whole. This dominance also raises a question of direction. *A. nerii* can itself alter milkweed chemistry, including cardenolide and latex production, and because we quantified cardenolides only at the end of the season, after a full season of aphid feeding, we cannot separate the possibility that host chemistry shaped aphid abundance from the possibility that sustained aphid pressure shaped the chemistry we measured. An experiment combining arthropod-exclusion treatments with repeated chemical sampling would separate these pathways and would allow the associations reported here to be tested causally.

### Conclusions

By tracking host chemistry, bacterial communities, and arthropods across four milkweed species, we tested how deterministic and stochastic processes contribute to phyllosphere community assembly across host species and through time. Our results show that the relative contributions of these processes were dynamic rather than fixed. Host identity was a major determinant of bacterial richness and composition, whereas inferred assembly processes shifted from early homogenizing dispersal toward increasing late-season dispersal limitation, with host-specific homogeneous selection contributing only temporally restricted and inconsistent signals rather than a sustained late-season increase. In other words, communities on different plants became more alike early in the season and more distinct later. Young, sparsely colonized leaves drew colonists from a shared environmental pool delivered by wind, rain, and arthropod visitors, so neighboring plants converged. As leaves matured and resident populations established, arriving cells encountered occupied surfaces, and exchange among plants declined, allowing each community to drift along its own trajectory. That this divergence outweighed any sustained rise in host-specific selection suggests the late-season imprint of host identity reflects the accumulated history of which taxa arrived first, as much as ongoing chemical filtering. Arthropods were associated with bacterial diversity in strongly host-dependent ways, further emphasizing that biotic interactions cannot be separated from host context. The broader implication is that phyllosphere community assembly cannot be adequately represented by a single temporal snapshot, *i.e.*, the relative contributions of dispersal, selection, host identity, and associated arthropods change over the growing season and differ among host species. Our findings therefore demonstrate the value of sampling repeatedly through a season, across trophic levels, and combining this with null-model approaches to investigate the dynamic assembly of plant-associated microbiomes. Future work should resolve the links between specific cardenolide compounds and bacterial taxa, the consequences of these community shifts for plant defense and growth, and whether the seasonal patterns observed here hold across years and locations.

## ACKNOWLEDGEMENTS

We thank Mary Arlin Bunnell, Hannah Thompson, Abigail Myers, and Zach Leopold for their help with setting up the experiment and data collection. We also thank Anurag Agrawal for reviewing the manuscript and providing valuable suggestions that helped improve the paper. This work was supported by the Department of Entomology at the Pennsylvania State University, the Department of Biology at the University of Alabama, the National Science Foundation, Division of Environmental Biology award #2440876, and the USDA National Institute of Food and Agriculture and Hatch Appropriations under Project #PEN04923, and Accession #7006440.

## DATA ACCESSIBILITY

1. The dataset and scripts can be found on Dryad platform. Link: <u>to be submitted</u>
2. The raw DNA sequence can be found on NCBI under the ID project: <u>to be submitted</u>

## STATEMENT OF AUTHORSHIP

MFK-B & VMP: conceptualized the research; MFK-B & VMP: conducted the experiment and collected the data; DFBS, GCB, carried out the sequencing of the bacterial DNA and bioinformatics; DFBS, MKB, and JGA worked on the cardenolide analysis. MFK-B, DFBS, LB, FD-A: analyzed the data; MFK-B and DFBS wrote the paper and all authors contributed substantially to revisions.

